# Zika virus NS3 drives the assembly of a viroplasm-like structure

**DOI:** 10.1101/2024.09.16.613201

**Authors:** Tania Sultana, Chunfeng Zheng, Garret Morton, Timothy L. Megraw

## Abstract

Zika virus (ZIKV) is a mosquito-transmitted flavivirus that caused an epidemic in 2015-2016 in the Americas and raised serious global health concerns due to its association with congenital brain developmental defects in infected pregnancies. Upon infection, ZIKV assembles virus particles in a virus-generated toroidal compartment next to the nucleus called the replication factory, or viroplasm, which forms by remodeling the host cell endoplasmic reticulum (ER). How the viral proteins control viroplasm assembly remains unknown. Here we show that the ZIKV non-structural protein 3 (NS3) is sufficient to drive the assembly of a viroplasm-like structure (VLS) in human cells. NS3 encodes a dual-function protease and RNA helicase. The VLS is similar to the ZIKV viroplasm in its assembly near centrosomes at the nuclear periphery, its deformation of the nuclear membrane, its recruitment of ER, Golgi, and dsRNA, and its association with microtubules at its surface. While sufficient to generate a VLS, NS3 is less efficient in several aspects compared to viroplasm formation upon ZIKV infection. We further show that the helicase domain and not the protease domain is required for optimal VLS assembly and dsRNA recruitment. Overall, this work advances our understanding of the mechanism of viroplasm assembly by ZIKV and likely will extend to other flaviviruses.

**Importance:** The Zika virus replicates its genome and assembles virus particles in the cytoplasm within the replication organelle, a large virus-induced compartment also called the viroplasm. It does this in part by remodeling the endoplasmic reticulum. However, how the virus directs the host cell to assemble the viroplasm is mostly unknown. This study shows that Zika virus non-structural protein 3 (NS3) is sufficient to assemble a viroplasm-like structures, and indicates that NS3 has a central role in assembling the viroplasm. Understanding how the virus assembles the viroplasm compartment and NS3’s role in it should significantly advance our understanding of the cellular mechanisms of virus infection. This study aims to gain more understanding of the Zika virus and its viroplasm along with the molecular mechanisms for viroplasm assembly which might be shared by other viruses.

## Introduction

The epidemic outbreak of the Zika virus (ZIKV) in 2015-2016 in the Americas raised serious health concerns worldwide due to the detrimental effects on fetal brain development and associated Guillain-Barré syndrome (GBS) in adults (1). ZIKV RNA has been detected in the placental and fetal brain tissues of infected pregnant mothers. Moreover, ZIKV can infect neurons and neuronal progenitor cells, leading to microcephaly and other brain developmental defects (2, 3).

ZIKV is a mosquito-transmitted flavivirus related to Dengue virus (DENV), yellow fever, West Nile, and Japanese encephalitis viruses (JEV) (4–6). The ZIKV genome is a positive-sense single-stranded RNA of approximately 11,000 bases in length that encodes three structural proteins (envelope E, prM/M, and capsid C) and seven non-structural proteins (NS1, NS2A, NS2B, NS3, NS4A, NS4B, and NS5) expressed as a single polyprotein that is processed by host and viral proteases into the 10 individual viral proteins (Fig. 1A) (7).

**Figure 1.**
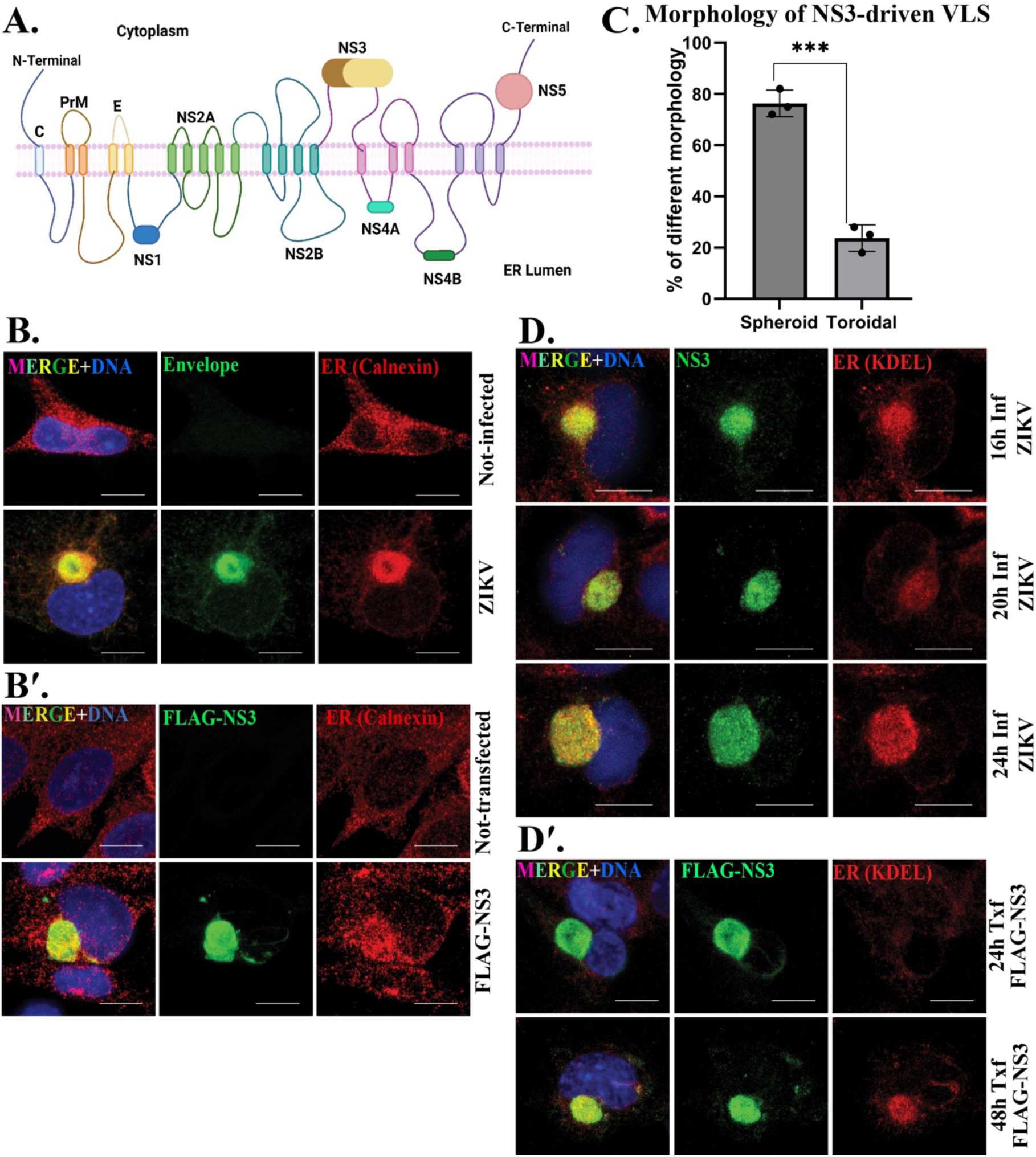
ZIKV NS3 drives assembly of a viroplasm like structure (VLS) which precedes ER recruitment. **(A)** Schematic of the ZIKV polyprotein and its topology within the ER membrane. Created with Biorender.com. Immunofluorescent (IF) staining of **(B)** mock and ZIKV-infected cells 24h post-infection (p.i) (viroplasm marked by ZIKV envelope protein; green) and (**B**′) mock and FLAG-NS3 transfected for 48h (NS3-driven VLS marked by FLAG; green) SNB19 cells. ER is marked with calnexin (red). **(C)** Morphology of the NS3-driven VLS. ER recruitment (**D**) to the viroplasm, marked with NS3, 16h (upper panel), 20h (middle panel), 24h (bottom panel) p.i and to the VLS (**D**′) 24h after transfection (upper panel) and 48h (bottom panel). ER is marked with KDEL (red). DAPI was used to stain the nucleus (blue) for IF staining in all figures. Scale bars: 10 μm.

Many viruses, including flaviviruses such as ZIKV, organize a compartment within the host cell refered to as a replication organelle, viroplasm, replication compartment, etc. from viral and host proteins and by reorganizing host organelles (8–12). The viroplasm is a distinct cytoplasmic compartment where virus replication, protein synthesis, and virus assembly occur. Its organization can vary among different types of viruses. Although the size of the viroplasm varies depending on the types of flavivirus and host cells, the flavivirus viroplasm is always derived from the endoplasmic reticulum (ER) (13–17). However, how the compartment assembles remains unclear. Consequently, gaining insights into ZIKV viroplasm formation and identifying the proteins that orchestrate its assembly will advance our understanding of the ZIKV infection cycle.

We previously showed that during ZIKV infection the viroplasm organizes around the centrosome and has a toroidal shape with the centrosome positioned within the core (18). In the absence of centrosomes, the viroplasm assembles but is spherical (18). Microtubules are required for efficient viroplasm assembly and the viroplasm reorganizes microtubules on its surface (18). Here, in our pursuit of understanding the roles of individual ZIKV proteins in viroplasm formation, we discovered a key role for the ZIKV NS3 protein. We found that ZIKV NS3 drives the assembly of a similar structure, a ‘viroplasm-like structure’ or VLS. ZIKV NS3 is a dual-domain protein with an N-terminal trypsin-like serine protease domain and a C-terminal helicase domain. The NS3 helicase utilizes the chemical energy derived from ATP hydrolysis to unwind double-strand (ds) RNA during replication (7) and contains functional motifs found in RNA helicases, including walker A and B motifs for ATP binding and hydrolysis, and a separate RNA binding domain (19). We further show that the assembly of the VLS by NS3 is impeded by mutation of the ATPase motifs but not by mutations in the protease or RNA binding domains.

## Results

### ZIKV NS3 is sufficient to form a Viroplasm-Like Structure (VLS)

The ER is typically a dispersed organelle in cells (Fig. 1B). However, in ZIKV-infected SNB-19 cells, it reorganizes and is incorporated into a spherical structure adjacent to the nucleus as it becomes integrated with the viroplasm (Fig. 1B).

We transfected plasmids that express each individual protein coding sequence of ZIKV strain MR766 with an N-terminal FLAG tag in SNB-19 cells, a human astrocytoma cell line, and found that NS3 was sufficient to generate a spherical compartment similar to the viroplasm (Fig. 1B, B′). None of the other ZIKV proteins generated such a structure when expressed from plasmids in cells (not shown). We refer to this structure as the viroplasm-like structure, or VLS, as it recruits the ER and results in nuclear deformation as does the virus-induced viroplasm (Fig. 1B, B′).

ZIKV-induced viroplasms assemble around the centrosome and have a toroidal shape 96% of the time (18). In contrast, the VLS is toroidal only 24% of the time, and is spherical with no discernable hollow core at a frequency of 76%, (Fig. 1C).

### NS3-driven VLS assembly precedes ER recruitment

We observed that ER recruitment to the VLS was variable 24 hrs after transfection, where a large compartment forms but is variable in the levels of ER recruited. Therefore, we conducted a comparative time-course analysis between the ZIKV-induced viroplasm and the NS3-driven VLS to delineate their distinct characteristics. The ZIKV-induced viroplasm shows no detectable delay in ER recruitment (Fig. 1D). However, we found that over the course of 24 h post transfection, NS3 expression produces VLS structures even though recruitment of the ER in the NS3-driven VLS was visibly weaker compared to 48h post transfection (Fig. 1D′). This highlights a distinct feature of the NS3-driven VLS where ER recruitment is delayed relative to the viroplasm where ER recruitment is immediate.

### NS3-driven VLS forms in association with the centrosome and recruits Golgi

The ZIKV viroplasm forms around the centrosome, and Golgi associates with the viroplasm surface (18). We investigated whether these associations were also features of the NS3-driven VLS and found that, similarly, the VLS forms in conjunction with the centrosome, although the localization of the centrosome was not predominantly positioned in the center of the compartment as it typically is with the viroplasm (Fig. 2A, B, C). The Golgi apparatus is typically organized into a single compact structure in SNB19 cells. In ZIKV-infected cells, the Golgi reorganizes into a more dispersed structure that is positioned on the surface and/or at the viroplasm core (Fig. 2D) (18). We observed a similar localization of the Golgi on the surface of the NS3-driven VLS and, less frequently, at the core of the VLS (Fig. 2 D′).

**Figure 2.**
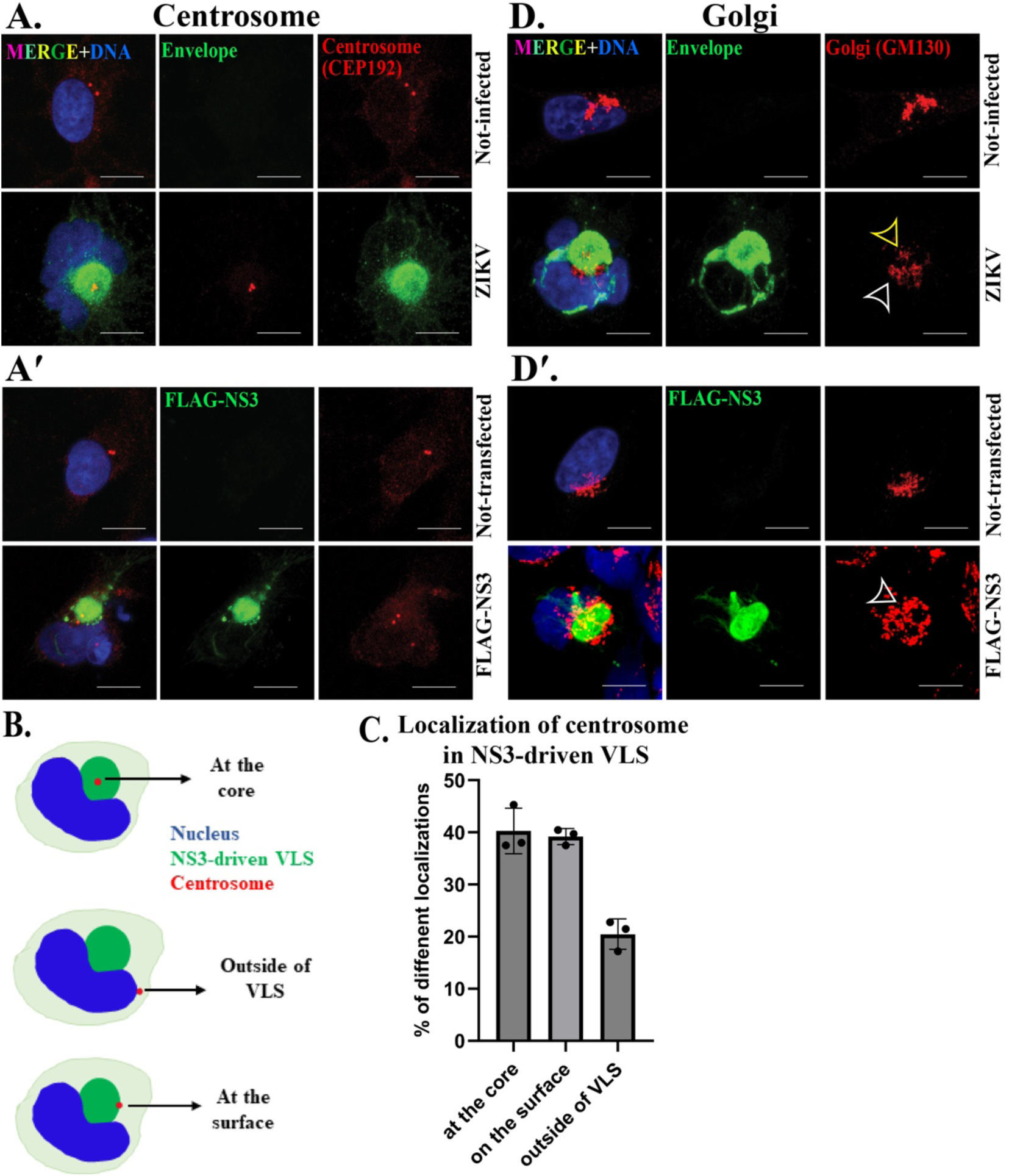
NS3-driven VLS forms in association with the centrosome and recruits the Golgi apparatus. IF staining for the centrosome protein CEP192 (red) in **(A)** mock and ZIKV (MR766) 24h post-infection and (**A**′) mock and FLAG-NS3 transfected for 48h in SNB19 cells. ZIKV-induced viroplasm (envelope protein, green) and NS3-driven VLS (FLAG, green). Illustration **(B)** and bar graph (mean ± standard deviation) shows different localization of centrosome in NS3-driven VLS. IF staining of the Golgi protein GM130 (red) in **(D)** mock and ZIKV infection for 24h and (**D**′) mock and FLAG-NS3 transfected for 48h in SNB-19 cells. The Golgi at the core is indicated by yellow arrowheads and the Golgi surrounding the viroplasm by white arrowheads. Scale bars: 10 μm.

### dsRNA localizes at the NS3-driven VLS

The ZIKV genome synthesizes dsRNA as a replication intermediate that localizes within punctate regions within the viroplasm (Fig. 3A) (18, 20). We produced fluorescent (fluorescein)-labeled dsRNA *in vitro* and co-transfected it into cells with the FLAG-NS3 plasmid to examine its localization relationship with the VLS. The dsRNA localized to the VLS, but did not display the similar punctate pattern characteristic of viral dsRNA at the viroplasm upon virus infection (Fig. 3. Thus, although qualitatively different, the VLS was capable of incorporating dsRNA.

**Figure 3.**
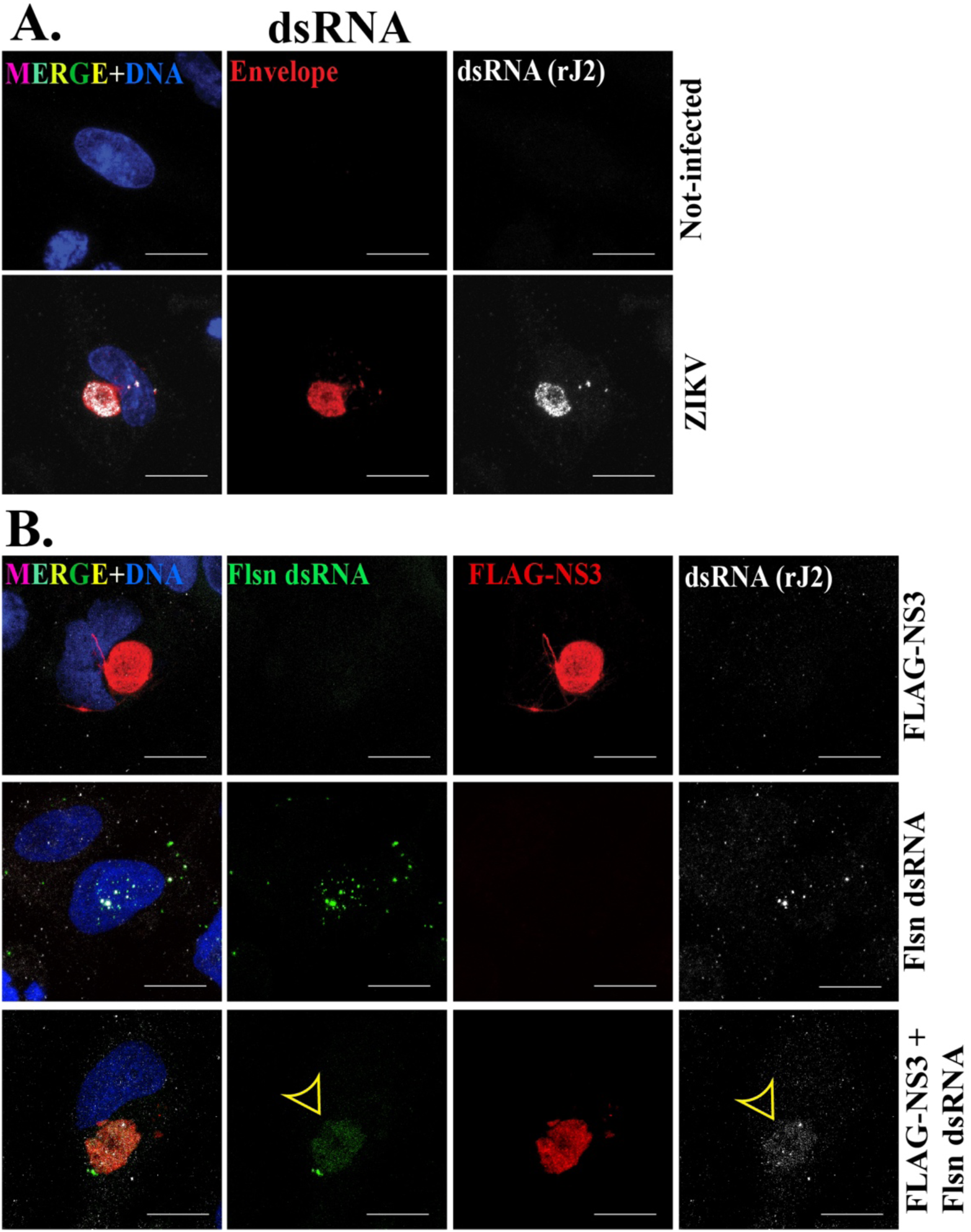
The NS3-driven VLS recruits dsRNA. IF staining of dsRNA (rJ2 antibody, white) **(A)** in mock and ZIKV (MR766) 24h post-infection. ZIKV-induced viroplasm is marked by ZIKV envelope protein; red. **(B)** FLAG-NS3 transfection (top panel) and fluorescein-labeled (Flsn) dsRNA transfection (middle panel) and co-trasnfection of Flsn dsRNA with FLAG-NS3 (bottom panel) for 48h in SNB19 cells. dsRNA recruitment is indicated by the yellow arrowheads. NS3-driven VLS is marked with FLAG (red). Scale bars: 10 μm.

### Microtubules are reorganized at the NS3-driven VLS

The extensive microtubule array in mock-infected cells undergoes reorganization upon virus infection, adopting a cage-like formation surrounding the viroplasm (Fig. 4A) (18). The reorganization of microtubules that takes place concurrently with viroplasm assembly prompted us to explore the arrangement of microtubules in the NS3-driven VLS. We discovered that microtubules similarly reorganized at the surface of the NS3-driven VLS, encircling its surface similarly to the viroplasm (Fig. 4B).

**Figure 4.**
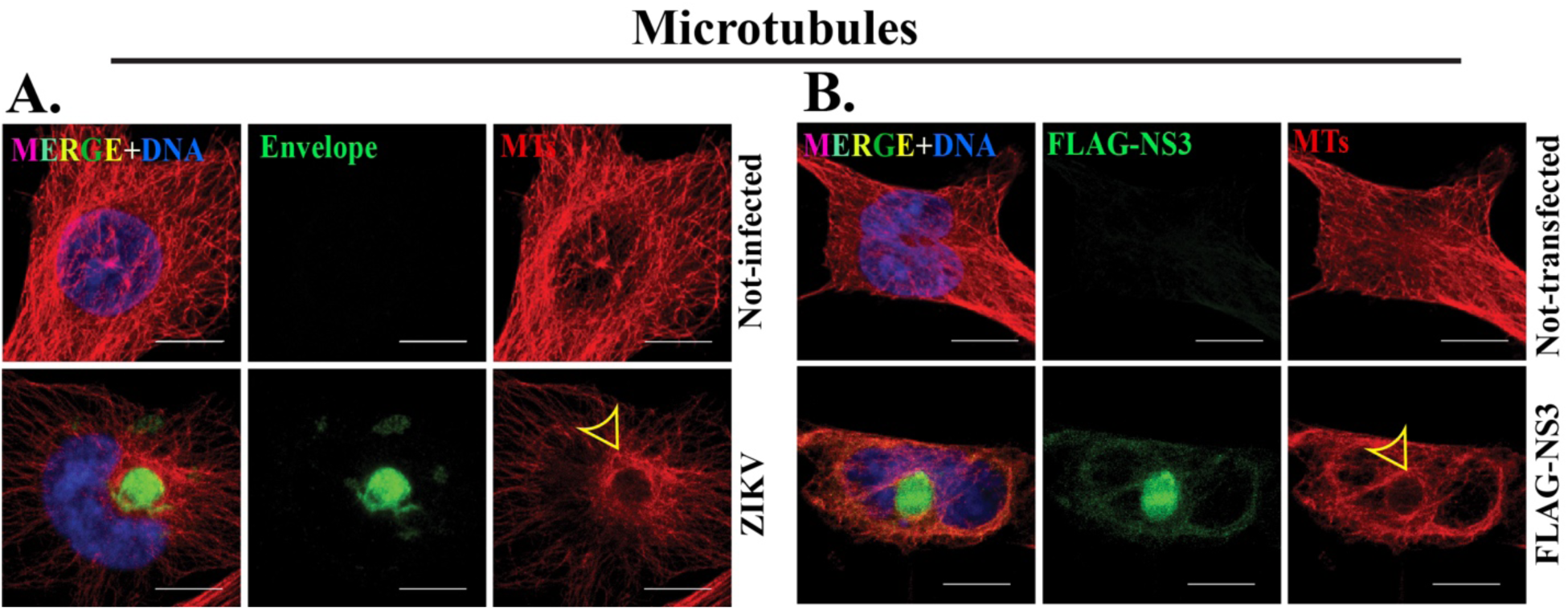
Microtubules are reorganized at the NS3-driven VLS. IF staining of microtubules (MTs) marked by α-tubulin (red) in **(A)** mock and ZIKV 24h post-infection and (**B**) mock and FLAG-NS3 transfected for 48h in SNB19 cells. ZIKV-induced viroplasm (green) and NS3-driven VLS (green) are marked with Envelope and FLAG respectively. Yellow arrowheads indicate MTs arrangement around the viroplasm and NS3-driven VLS. Scale bars: 10 μm.

### VLS formation by NS3 requires helicase activity

Having demonstrated the sufficiency of NS3 to drive VLS assembly, we next tested which domains of NS3 are required. We created mutations in functional residues of the protease or helicase domains of NS3 (21–23) such as serine to alanine (S135A) in the protease domain, and for ATP binding and hydrolysis, lysine to asparagine (K210N) in the walker A motif, and aspartic acid to asparagine (D290N) in the walker B motif, respectively, and arginine to glutamine (R461Q) essential for nucleic acid binding in the helicase domain (Fig. 5A). See a summary of the mutants in Table 2. All of the mutant proteins expressed at similar levels to the wild-type NS3 protein when expressed in transfected cells (Fig. 5B). We then investigated the impact of these mutations on VLS formation.

**Figure 5.**
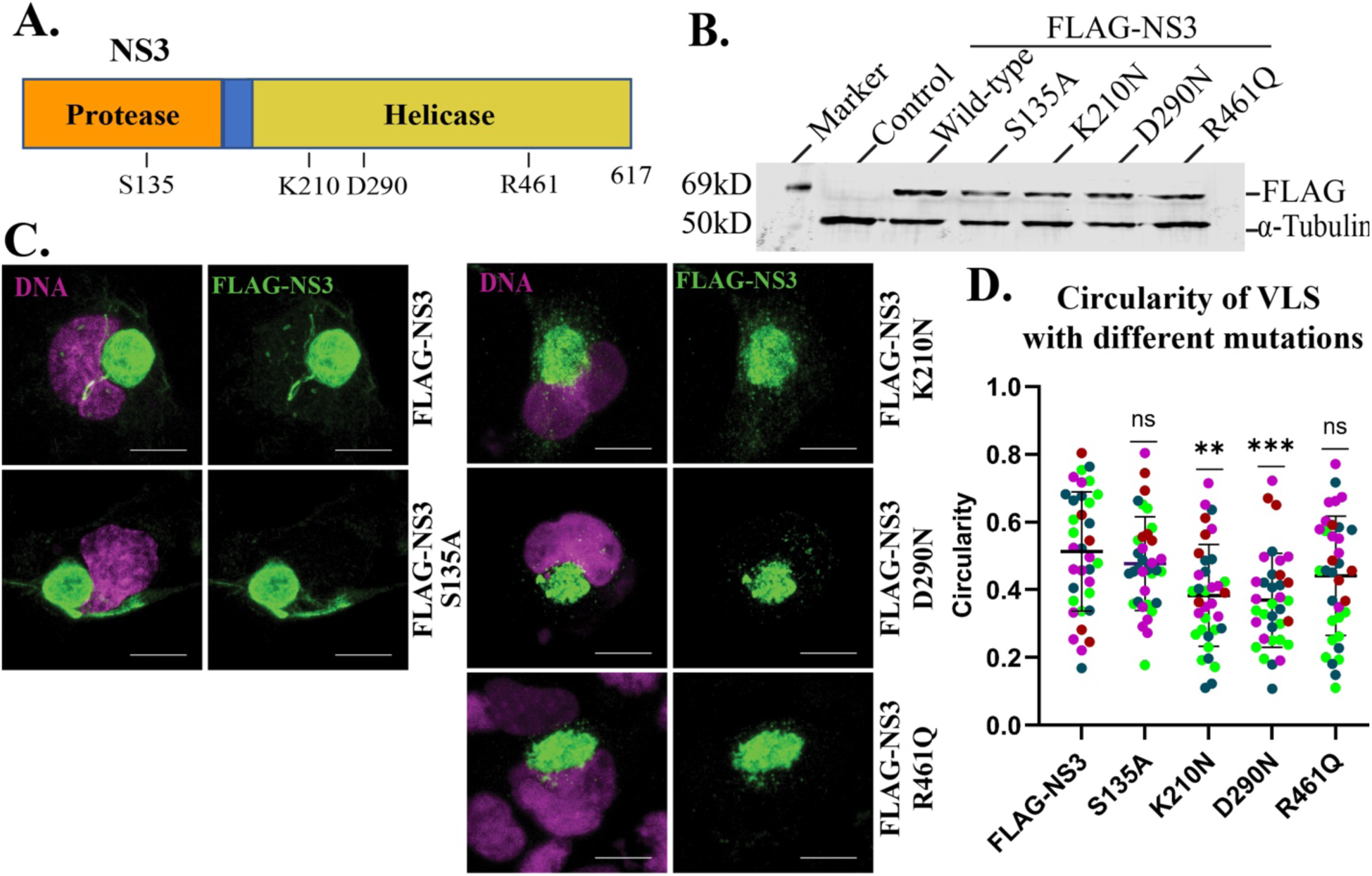
VLS formation by NS3 requires the ATPase domain. **(A)** NS3 structure denoting protease and helicase domains and positions of introduced mutations. **(B)** Western blot of FLAG-tagged wild-type and mutant NS3 proteins. **(C)** IF staining of FLAG-NS3 and mutated NS3 (S135A, K210N, D290N and R461Q) transfected for 48h in SNB19 cells. NS3-driven VLS is marked with FLAG (green). **(D)** Circularity of VLS with different mutations. Different colors indicate different experiments (n=4). \**P* < 0.05, ** *P* < 0.001, and \*\*\**P* < 0.0001 are considered statistically significant differences from the control (FLAG-NS3). DAPI stains the nucleus (magenta). Scale bars: 10 μm.

**Table 1.**
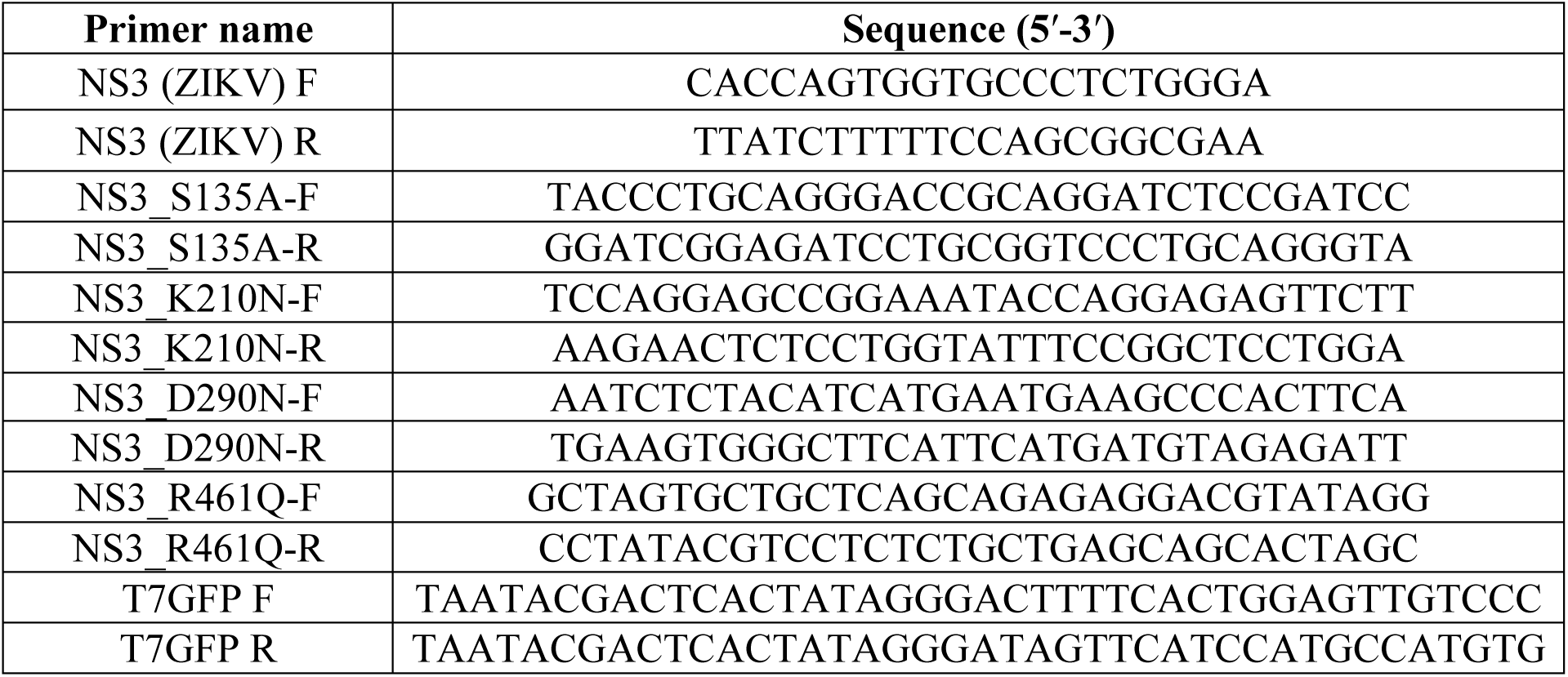
Primers for cloning and mutagenesis of NS3 and for dsRNA template.

**Table 2:**
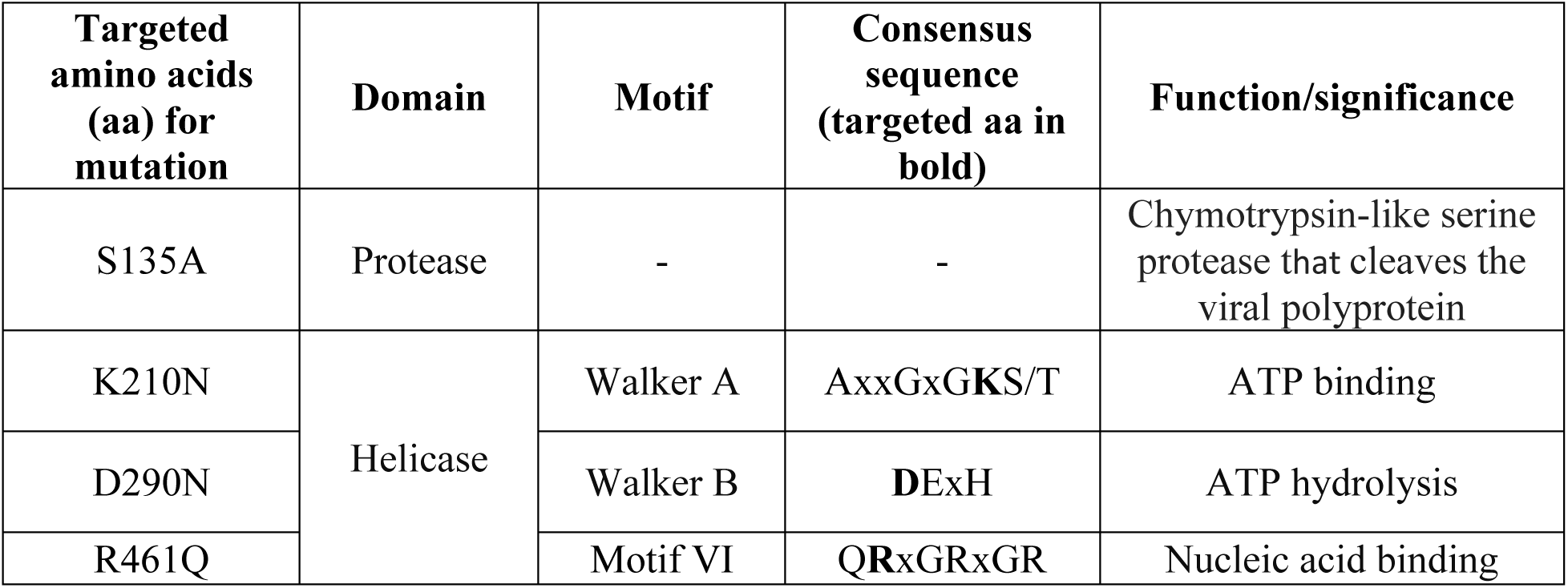
Summary of mutations generated in NS3.

All of the mutant proteins were capable of generating a VLS to some degree, but the mutations in the ATPase Walker A and B motifs (K210N and D290N, respectively) produced less organized VLS structures that were less compact (Fig. 5C, D). Mutations in the protease and RNA-binding domains, on the other hand, resulted in no detectable impairment in the ability of NS3 to assemble the VLS (Fig. 5C, D).

## Discussion

Here we show that the ZIKV NS3 protein is sufficient to assemble a compartment within human cells with the features of a viroplasm. With multiple features similar to the viroplasm that forms at virus infection, we refer to the NS3-generated compartment as a viroplasm-like structure (VLS). The VLS has multiple features that relate it to the ZIKV viroplasm including the recruitment of ER, Golgi, and dsRNA, its organization in association with the centrosome, and the organization of microtubules at its surface (18).

During ZIKV infection the mature viroplasm typically exhibits a toroidal (ring-shaped) morphology, and a spherical shape less frequently. This viroplasm morphology is also seen with a Puerto Rican strain of ZIKV and with DENV serotype 2 (18). However, the NS3-driven VLS displays a toroidal shape less frequently than the viroplasm (Fig. 1C). NS3 alone may initiate VLS formation but might lack the additional viral and cellular machinery to fully replicate the architecture of the mature viroplasm. Previous investigations have also shown co-localization of flavivirus NS3 with the ER (24–26). However, in contrast with the dynamics of viroplasm formation during ZIKV infection, the VLS appears capable of forming prior to ER recruitment (Fig. 1D, D′) indicating that ER recruitment is not limiting for VLS assembly. Several factors could contribute to the variability of ER localization/reorganization at the VLS. One possibility is that the process of VLS formation may be slow, relative to the viroplasm, to fully engage and integrate the ER. In addition, in the viral replication process the ZIKV genome is expressed as a polyprotein within the ER and so native NS3 is expressed and processed at the ER in contrast with the FLAG-NS3 protein we expressed from a plasmid. This could partially explain the delay in VLS assembly and ER recruitment relative to the viroplasm. Moreover, while these data indicate that NS3 is sufficient to generate a VLS and recruit ER, other ZIKV proteins might be needed for full efficiency.

In ZIKV infected cells, the centrosome typically lies at the core of the viroplasm and is associated with tyrosinated and acetylated microtubules (18). However, the NS3-driven VLS lacks the characteristic central location of the centrosome (Fig. 2A-C), Golgi localization (Fig. 2D, D′) and cluster of microtubules at the core (Fig. 4). One potential explanation for the distinct structural outcomes compared to the ZIKV-induced viroplasm could be variations in the assembly kinetics or a lack of accessory ZIKV factors to facilitate VLS assembly In addition to its enzymatic functions, NS3 likely engages in interactions with host proteins, contributing to the subcellular rearrangements observed during ZIKV infection and with NS3 expression alone. Previous research indicates that NS3 interacts with several host proteins, including those associated with innate immunity, lipid biosynthesis, ER and Golgi trafficking, and the cytoskeleton (24–35). More investigation will be required to fully understand how NS3 accomplishes assembly of the compartment. Nonetheless, the mutation analysis we conducted indicates that the ATP binding and/or ATPase activity of NS3 is involved.

The centrosome serves as the primary microtubule-organizing center (MTOC) in proliferating animal cells. In addition to the centrosome, other non-centrosomal MTOCs (ncMTOCs) such as the Golgi apparatus organize microtubules at numerous locations within the cell to support a variety of cellular functions (36–43). ZIKV infection disrupted centrosome organization and mitotic irregularities in neural progenitors, resulting in altered neural progenitor differentiation (44–46). Moreover, ZIKV NS3 was found to be associated with the centrosome and several centrosomal proteins (27, 28, 30, 31, 47). In addition, expression of NS3 was sufficient to modify centrosome architecture and alter the localization of the centrosome protein CEP63, consequently suppressing the innate immune response (44). In our work we did not detect NS3 localization at centrosomes, an indication that that localization might be cell-type specific.

However, the strong accumulation of NS3 at the viroplasm and VLS in association with the centrosome might mask detection of binding there. The VLS formed in association with the centrosome (Fig. 2A, A′) and Golgi apparatus with the centrosome positioned in the center or on the periphery of the VLS, while the Golgi was associated with the VLS surface (Fig. 2D, D′). This contrasts with ZIKV-infected cells, where the centrosome and some Golgi were more often found at the core, but which is less frequent with the VLS.

ZIKV RNA synthesis initiates with the production of negative-sense RNA which results in a dsRNA intermediate (Fig. 3A). New positive-sense RNA genome copies are generated from these dsRNA templates (48). We observed the localization of exogenous dsRNA at the NS3-driven VLS (Fig. 3B). However, the pattern of localization is strikingly different between the viroplasm, with punctate sites of dsRNA localization, compared to the VLS with dsRNA localized throughout the structure. This difference is unclear, but might be related to the mechanics whereby the generation of dsRNA in the viroplasm are generated *in situ*, whereas in the experimental approach we took dsRNA was exogenous and introduced pre-formed.

Many different viruses rely on microtubules and alter microtubules to facilitate the intracellular trafficking of virus components (49–57) including flaviviruses (15, 18, 58–61). Microtubule arrays were shown to be localized at the mature viroplasms of various flaviviruses such as DENV, Kunjin virus, ZIKV and tick-borne encephalitis virus (15, 18, 62–64). NS3 of JEV, ZIKV and Kunjin virus was found to be associated with microtubules (24, 47, 63). It was predicted that NS3 might induce microtubule reorganization in JEV-infected cells in order to facilitate the transport of other viral proteins from one organelle to another during JEV replication (24). Our previous findings along with other investigations reveal that microtubules form a cage-like arrangement in close proximity of mature viroplasm in ZIKV-infected cells (15, 18). As was shown with the viroplasm upon ZIKV infection, the arrangement of microtubules in the VLS surrounded the VLS in a cage-like structure (Fig. 4A, B,). The organization of microtubules at the surface of the VLS indicates that NS3 is sufficient for the recruitment of factors that control microtubule nucleation, stability or anchoring at the surface of the VLS. Additional studies will be necessary to understand the host factors that get recruited to the VLS to impart the organization of microtubules there similar to how they are organized at the viroplasm (18). Earlier reports discovered the association of JEV and ZIKV NS3 with microtubules (24, 47), indicating that this could be a conserved role among flavivirus NS3 proteins.

To understand further the mechanisms by which NS3 can organize the VLS compartment, we expressed mutant versions of NS3 with mutations that should disrupt the protease, helicase ATPase, and the RNA binding domains (21, 22). While previous work indicated the protease and helicase were interdependent for optimal functionality, we found that only mutations that disrupted the ATPase walker A or walker B motifs impaired organization of the VLS by NS3 (Fig. 5C, D). Overall, these findings indicate that the ATPase activity of NS3 is required for VLS formation.

In conclusion, we report that ZIKV NS3 is sufficient to form a VLS that recruits the ER, reorganizes Golgi to its surface, forms in association with the centrosome, and organizes microtubules at its surface. Altogether, the VLS mirrors the viroplasm that is formed upon ZIKV infection. We further show that the ATPase subdomain in the helicase domain is required for efficient VLS formation. Overall, these findings indicate that NS3 plays a key role in viroplasm assembly during ZIKV infection. The design of inhibitors to flavivirus NS3 might therefore be effective in impeding the virus in more ways besides inhibiting RNA replication.

## Materials and Methods

### Cell culture

We used the U-251 cell line derivative SNB19 cells (Charles River Laboratories, Inc. under contract of the Biological Testing Branch of the National Cancer Institute) for all virus infections and plasmid transfections. SNB19 cells were cultured and maintained in RPMI medium (Cytiva-HyClone, cat# SH30027.02) on standard cell culture plates. Virus stock production was carried out using Vero E6 cells (ATCC) cultured in DMEM media. All media were supplemented with 10% fetal bovine serum (FBS) (Avantor Seradigm, cat# 97068-85) and 1% penicillin-streptomycin (Corning, cat# 30-002-CI) and maintained in a humidified environment with 5% CO_2_ at 37°C.

### Plasmids and reagents

We employed Gateway cloning (Invitrogen) to create entry clones in pENTR (Invitrogen, cat#45-0218) and subsequently subcloned inserts into the destination vector, pSG5-FLAG (a gift from Eric C. Johannsen (65), originally from Hatzivassiliou et al. (66) by recombination using LR Clonase (Invitrogen, cat#11791-020). All ZIKV NS3 sequences were from ZIKV strain MR766. The primers for cloning NS3 constructs are listed in Table 1.

### Transfection

Cells were plated for ∼24h in 12-well plates (Greiner, cat # 665180) containing sterile 18 mm circular coverslips (Electron Microscopy Sciences, cat#72222-01). Upon reaching 70%−80% confluency, cells were transfected using 1ug of plasmid for 24h for the detection of early ER recruitment and 48h for all other experiments. Lipofectamine 3000 reagent (Invitrogen, cat# L3000015) was used following the manufacturer’s protocol for 12-well plates. After transfection, the media was removed and cells were fixed with 100% methanol (VWR Chemicals, cat# BDH2018-1GLP) at −20°C for 10 minutes followed by three phosphate-buffered saline (PBS) washes. Untransfected cells were used as control.

### Virus Stock Production and Infection

The ZIKV-MR766 strain was obtained from ZeptoMetrix. Virus stocks were prepared as described previously (67). In brief, we infected Vero E6 cells with virus stock diluted in the culture medium at a multiplicity of infection (MOI) of 0.01 and allowed them to incubate with the cells for 2 hours. The infection medium was replaced with fresh media, and the supernatant was harvested 72–96 hours post-infection (p.i.) when a significant cytopathic effect was observed in the majority of the cells. The supernatants were centrifuged at 1000 x g for 10 minutes, filtered through a 0.45-micron filter (VWR Chemicals, cat# 76479-028), aliquoted, and stored at −80°C.

SNB19 cells were infected using diluted virus in culture medium at a MOI of 1.0 for 2 hours. Subsequently, we replaced the viral media with fresh culture media and maintained it for 24 hr. Cell culture media was used for mock infection.

### Synthesis of Fluorescein labeled dsRNA

dsRNA was synthesized following the previously described method (68). Briefly, GFP coding sequence was amplified by PCR and used as a template for in-vitro transcription with T7 polymerase. Table 1 lists the sequences of the primers used for PCR. HyperScribeTM T7 High Yield Fluorescein RNA Labeling Kit (APExBIO, cat# K1060) was utilized for dsRNA synthesis. The RNA was purified using phenol-chloroform extraction followed by ethanol precipitation. The concentration and purity of the RNA were measured using a NanoDrop spectrophotometer. 0.25ug of RNA was used for co-transfection with pSG5-FLAG-NS3.

### Site-directed Mutagenesis

The pSG5-FLAG-NS3 plasmid was used as the target for site-directed mutagenesis. Mutants S135A, K210N, D290N and R461Q were constructed using QuikChange II site-directed mutagenesis kit (Agilent, cat# 200523-5) according to the manufacturer’s protocol. Subsequently, mutations were confirmed by DNA sequencing. Table 1 lists the sequences of the primers used for site-directed mutagenesis.

### Western Blotting

SNB-19 cells were seeded in 6 well plates (Greiner, cat# 657160) followed by transfection with wild-type and mutant plasmids. After 24h, cells were washed with cold PBS, harvested, and lysed in SDS-PAGE loading dye. The lysates were centrifuged at 4°C for 15 minutes, and the supernatants were boiled at 95°C for 5 minutes before loading onto SDS-PAGE gels. Protein bands were transferred to nitrocellulose membranes (VWR, cat# 28298-020) which were blocked with 5% non-fat milk in Tris-Buffered Saline with Tween 20 (TBST). Following washing, membranes were incubated with primary antibodies overnight at 4°C and then with secondary antibodies for 2 hours at room temperature (RT). Protein bands were visualized using a LiCor Odyssey imaging system and analyzed using Image Studio Ver 5.2 (LiCor Biosciences).

### Immunofluorescent (IF) Staining and Microscopy

For IF staining, MeOH-fixed cells were treated in primary antibody in PBS solution containing 0.1% saponin (Sigma-Aldrich, cat# SAE0073-10G) and 5 mg/mL BSA for 2h at RT or overnight at 4°C, washed three times for 5 min in PBS, and then incubated with secondary antibodies for 2 h at RT. The wells were washed three times for 5 min with PBS and then mounted on slides in mounting medium (68).

Slides were imaged on a Nikon AX confocal microscope with a 60×NA 1.49 oil immersion objective using NIS-elements (Nikon) software. Region of interest (ROI) in ImageJ software was used for circularity measurements (69).

### Antibodies

The following primary antibodies were used for staining: chicken monoclonal FLAG (Exalpha Biologicals, Inc cat# AFLAG, 1:500 for IF), rat monoclonal FLAG (Agilent, cat# 200473, RRID: AB_10596510, 1:1000 for IF, 1:10000 for WB), rabbit polyclonal ZIKV envelope protein (Kerafast, cat# EFS001, 1:500 for IF), mouse monoclonal anti-flavivirus group antigen (envelope) (EMD Millipore, D1-4G2-4-15, cat# MAB10216, 1:1000 for IF), mouse monoclonal anti-KDEL (Enzo Life Sciences, 10C3, cat# ADI-SPA-827, RRID: AB_2039327, 1:500 for IF), rabbit monoclonal anti-calnexin (Cell Signaling Technology, C5C9, cat# 2679S, RRID:AB_2228381, 1:1000 for IF), and rabbit monoclonal anti-GM130 (Cell Signaling, D6B1, cat #12480, RRID: AB_2797933 1:3000 for IF), mouse monoclonal anti dsRNA, rJ2 (EMD Millipore, cat# MABE1134, RRID: AB_2819101, 1:500 for IF), mouse monoclonal anti-alpha-tubulin (DM1A, Sigma-Aldrich, cat# T9026, 1:1000 for IF) and mouse monoclonal anti-GCP2 (Clone 01, a gift from Pavel Dráber, 1:500 for IF) (70).

We used goat anti-mouse secondary antibody (LI-COR, IRDye 800CW, cat# 926-32210, RRID:AB_621842) 1:20,000 for WB and Alexa Fluor 488, 568, and 647 conjugated goat secondary antibodies (Invitrogen, 1:1000) for IF staining and the cell nuclei were stained using DAPI (Invitrogen, cat# D1306, 1ug/ml for IF).

### Statistical Analysis

The results in all graphs are presented as the mean ± standard deviation (SD) and analyzed by GraphPad Prism version 10.0.0 for Windows (GraphPad Software, Boston, Massachusetts USA). An unpaired Student’s t-test was carried out to determine the differences in means between experiments with a probability level of \**P* < 0.05, ** *P* < 0.001, and \*\*\**P* < 0.0001 considered increasingly significant, respectively.

## Acknowledgements

We appreciate the Megraw lab members for their insightful discussions and feedback on the manuscript. We also thank Dr. Fanxiu Zhu, professor of Biological Sciences, Florida State University for his insights and feedback on the project. We thank Leanne Duke, a graduate student in the Meckes Lab at FSU for assistance with cell culture and virus infection. We appreciate Rebecca A. Buchwalter, a former graduate student in the Megraw lab, for generating the preliminary findings for this project.

## Data availability

The data presented in this study is available upon request from the corresponding author.

## Author’s contributions

T.L.M. conceived the project. T.S. performed experiments, analyzed data, and wrote the draft manuscript. C.F. assisted in experiments, study design and manuscript revision. T.L.M secured funding, supervised and revised the manuscript. All authors have read and agreed to the published version of the manuscript.

---

Table 1 lists the sequence of the primers and Table 2 shows the functions of selected amino acids in protease and helicase domain.

